# A snakemake toolkit for the batch assembly, annotation, and phylogenetic analysis of mitochondrial genomes and ribosomal genes from genome skims of museum collections

**DOI:** 10.1101/2023.08.11.552985

**Authors:** Oliver White, Andie Hall, Ben W. Price, Suzanne T. Williams, Matt Clark

## Abstract

Low coverage “genome-skims” are often used to assemble organelle genomes and ribosomal gene sequences for cost effective phylogenetic and barcoding studies. Natural history collections hold invaluable biological information, yet poor preservation resulting in degraded DNA often hinders PCR based analyses. However, with improvements to molecular techniques and sequencing technology, it is possible to use methods developed for working with ancient DNA to generate libraries and sequence the short fragments typical of degraded DNA to generate genome skims from museum collections.

Here we introduce a snakemake toolkit comprised of three pipelines *skim2mt*, *skim2rrna* and *gene2phylo*, designed to unlock the genomic potential of historical museum specimens using genome skimming. Specifically, *skim2mt* and *skim2rrna* perform the batch assembly, annotation and phylogenetic analysis of mitochondrial genomes and nuclear ribosomal genes, respectively, from low-coverage genome skims. The third pipeline *gene2phylo* takes a set of gene alignments and performs phylogenetic analysis of individual genes, partitioned analysis of concatenated alignments and a phylogenetic analysis based on gene trees.

We benchmark our pipelines with simulated data, followed by testing with a novel genome skimming dataset from both recent and historical solariellid gastropod samples. We show that the toolkit can recover mitochondrial and ribosomal genes from poorly preserved museum specimens of the gastropod family Solariellidae. In addition, the phylogenetic analysis of multiple gene sequences is consistent with our current understanding of taxonomic relationships.

The generation of bioinformatic pipelines that facilitate processing large quantities of sequence data from the vast repository of specimens held in natural history museum collections will greatly aid species discovery and exploration of biodiversity over time, ultimately aiding conservation efforts in the face of a changing planet.

## Introduction

Natural history collections are home to more than one billion, expert-verified specimens worldwide (Bartolozzi et al., 2023) as well as large numbers of unclassified or even bulk samples, and as such represent a vast repository of historical biological data that remains largely untapped. Challenges associated with such material include poor preservation, the use of unknown preservatives, ongoing DNA degradation and contamination. Such characteristics typically make direct recovery of intact gene sequences by polymerase chain reaction (PCR) impossible, without the amplification of many short overlapping fragments (D’Ercole et al., 2021). Fortunately, advances in novel laboratory techniques (Ruane & Austin, 2017; Straube et al., 2021) and next generation sequencing (NGS) technology make it possible to obtain DNA sequences from many historical specimens, unlocking the potential for wide-ranging genomic analyses. Using natural history collections provides the opportunity to easily work on many species, even if they are now extinct, rarely collected, or from areas of the world that are poorly sampled. Given that almost all known species have vouchers in one or more natural history collections, the use of these specimens could rapidly fill gaps in DNA reference libraries, greatly accelerating biodiversity discovery and DNA-based monitoring of the environment.

“Genome skimming” is a term referring to the generation of low coverage whole genome sequence data, first coined by Straub et al (2012). Although genome skimming does not generate data with sufficient coverage to assemble the entire nuclear genome, there are sufficient reads to assemble sequences that are present in the genome in multiple copies and are therefore highly represented in the sequence data. Common targets for genome skimming studies include organelle genomes (a cell has one nucleus but many organelles) and nuclear ribosomal genes (there are typically 100s or more copies of nuclear rRNA genes). Organelle and ribosomal genes have been widely used for phylogenetics due to their discriminatory power and the availability of “universal” primers and continue to be employed as “barcode” genes for identification. Specifically, the mitochondrial gene *cox1* for animals, chloroplast genes *matK* and *rbcL* for plants, 16S rRNA for bacteria and 18S or ITS for fungi are used now and dominate barcode sequence databases. In addition, novel genome skimming approaches are increasingly being developed, to use all sequence data to assign species identity via a “DNA-mark” (Bohmann et al., 2020) or “varKode” (De Medeiros et al., 2024).

When working with historical specimens in particular, genome skimming offers many advantages over polymerase chain reaction (PCR) amplification and Sanger sequencing of individual genes. Relatively long fragments of intact genomic DNA are required for PCR, whereas degraded and low amounts of DNA (∼1ng) typical of museum specimens (Mullin et al., 2023), are also suitable for short read NGS platforms. The wet lab work is relatively straightforward, only requiring DNA extraction and library methods optimised for degraded DNA which have been routinely used in the ancient DNA community for many years. Genome skimming also has additional benefits over targeted PCR since multiple loci can be recovered at the same time without development and optimisation of multiple PCR assays, or the need to design group specific DNA capture baits for target enrichment (Call et al., 2023). With advances in bioinformatic tools, it is likely that low coverage genome skimming datasets will have even greater utility in the future. For example, K-mer based approaches have been developed for genome skims to investigate phylogenetic relationships (Sarmashghi et al., 2017) and genome properties including genome length and repetitiveness (Sarmashghi et al., 2021). Finally, genome skimming is increasingly cost effective as the cost of NGS sequencing continues to decrease. In the light of these advantages, genome skimming is seen as a hugely scalable process that is suitable for batch recovery of useful genomic data from museum collections.

However, few bioinformatic pipelines are available to assist with the assembly of large numbers of organelle and nuclear ribosomal sequences from batches of genome skimming data. Notable exceptions include MitoZ (Meng et al., 2019) and NOVOWrap (Wu et al., 2021) for the assembly and annotation of mitochondrial genomes. MitoGeneExtractor can be used to extract mitochondrial protein coding genes from next generation sequencing datasets (Brasseur et al., 2023). In addition, plastaumatic (W. Chen et al., 2022) is available for chloroplast assembly and annotation and PhyloHerb (Cai et al., 2022) can be used for the assembly of chloroplast and nuclear ribosomal repeats without annotation. Other targeted assembly approaches are available including Orthoskim (Pouchon et al., 2022) for chloroplast, mitochondrial and ribosomal sequences. However, these tools were not designed with historical and/or degraded samples in mind nor issues such as contamination and the undesirable assembly of non-target sequences. In addition, these tools do not implement phylogenetic analysis of the annotated genes identified.

Here we introduce a snakemake toolkit comprised of three pipelines: *skim2mt*, *skim2rrna* and *gene2phylo*. These pipelines are designed to unlock the potential of historical museum specimens when using genome skimming. Specifically, *skim2mt* and *skim2rrna* perform the batch assembly, annotation and phylogenetic analysis of mitochondrial genomes and nuclear ribosomal genes, respectively. The third pipeline *gene2phylo* takes a set of gene alignments and performs phylogenetic analysis of individual genes, partitioned analysis of concatenated alignments and a phylogenetic analysis based on gene trees. The pipelines wrap 12 published bioinformatic tools as well as custom python and R scripts into a single user-friendly pipeline designed to cope with more challenging data from historical collections, permitting large scale genome skimming studies from museum specimens. These pipelines have many advantages for such studies, for example they (1) run on a single machine or in parallel on a High Performance Computing cluster, (2) process batches of samples, (3) assemble both mitochondrial and nuclear ribosomal sequences (4) use GetOrganelle (Jin et al., 2020) which an independent review found to be the best performing assembly tool for organelle genomes (Freudenthal et al., 2020), (5) perform basic assembly checking e.g. for contamination and non-target sequences (6) generate phylogenetic gene trees based on annotated genes. Whilst it would be possible to wrap all steps into one single pipeline, it is necessary to implement mitochondrial and ribosomal assembly individually, and user feedback suggested it was necessary to check outputs manually for evidence of contamination before more detailed phylogenetic analysis (see Result and Discussion for more details).

To benchmark the utility of our pipelines, we generated a simulated data for 25 species from Papilionoidea, a superfamily of butterflies with reference genomes and known taxonomy. We then analysed a novel genome skimming dataset for the gastropod family Solariellidae (hereafter solariellid gastropods). This group was selected as it represents many of the challenges associated with genome skimming museum collections. Solariellids are small marine snails found predominantly in deep-water. Many member species are rare, and as a family they are poorly represented in museum collections worldwide, with few live-collected specimens: many species are known only from a single, dry and often damaged shell (Williams et al., 2020). Although solariellid gastropods have been the focus of previous molecular phylogenetic studies (Sumner-Rooney et al., 2016; Williams et al., 2013, 2022), these studies have relied on partial sequence from only four genes, which have not fully resolved relationships among genera. As such, our understanding of solariellid evolution would greatly benefit from increasing the number of gene sequences used, furthermore there are no published reference genomes for the group with limited sequence data on public databases. Here we demonstrate how a genome skimming approach and our snakemake toolkit can be used to improve our understanding of phylogenetic relationships even in this challenging case.

## Material and methods

### Simulated data

Simulated data for mitochondrial and ribosomal sequences were generated for 25 species from the butterfly superfamily Papilionoidea. Simulated data were generated using gargammel (Renaud et al., 2017) with the following parameters: read length 150, location 5, scale 0.5, target coverage 20×, sequencing system HiSeq2500 and no contaminant sequences. A full list of taxa and reference datasets used to generate the simulated data are presented in Supporting Information Table S 1. Note that the simulated data and config files for this analysis are available with all pipelines as test data.

### Solariellid sample selection and sequencing

A total of 25 samples were selected, with representatives from 18 genera, encompassing the diversity of the solariellid family (Table 1). Samples differ in several ways that likely affected DNA quality and yield (Supporting Information Table S 2), for example, time since collection (1967-2015) and preservation method (dry shell with dehydrated body tissues or live-collected snail preserved in 70-99% ethanol). In addition, some shells were cracked, allowing the rapid penetration of ethanol, which is particularly important as snails can seal their bodies inside their shells by closing their operculum effectively excluding ethanol. Samples where shells have not been cracked generally have very degraded DNA. Samples also differ in time between sequencing and when DNA was extracted (0-10 years; Supporting Information Table S 2). DNA was isolated using Qiagen DNeasy blood and tissue kit and quantified using a Qubit fluorimeter and High Sensitivity assay kit. A Tapestation 2200 (Agilent Technologies, Santa Clara, USA) was also used to assess DNA integrity prior to library preparation. Polymerase Chain Reaction (PCR) amplification and Sanger sequencing of mitochondrial (*cox1*, 16S and 12S) and ribosomal genes (28S) were attempted for each sample at the time of DNA extraction and these results are compared with our genome skimming approach. Illumina Libraries were prepared using a SparQ DNA Frag and Library Prep kit (QuantaBio, Beverly, USA) and sparQ PureMag Beads (QuantaBio), with Sparq Adaptor Barcode sets A and B (QuantaBio). Libraries were normalised and pooled equally before being sent to Novogene (Cambridge, UK) for sequencing. The single indexed libraries were sequenced on an Illumina Novaseq on an S4 300 cycle flowcell using 150bp paired reads (see Data and code availability statement).

**Table 1.**
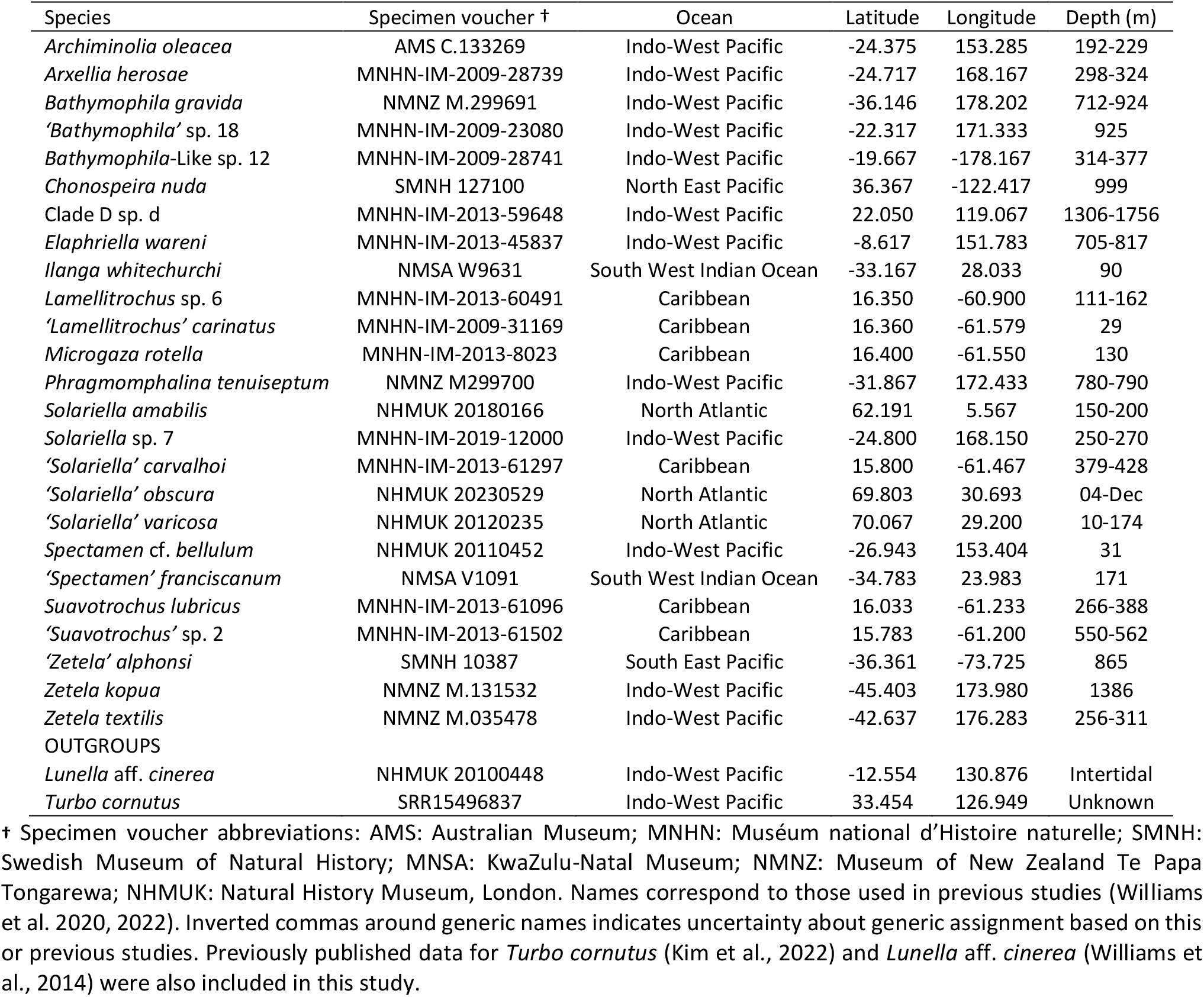
Sample details for 25 solariellid gastropod species and two outgroup species used in this study with museum registration numbers or NCBI sequence read archive number for sequence data (*Turbo cornutus* only), ocean of origin, latitude, longitude, and depth of collection location.

Additional sequence data for ‘*Solariella*’ *varicosa* were provided by Andrea Waeschenbach (Natural History Museum London, UK). Raw sequence data for two outgroups from the family Turbinidae were also analysed, including: *Turbo cornutus* (Kim et al., 2022; SRR15496837) and unpublished raw data for *Lunella* aff. *cinerea* (mitochondrial genome published in Williams et al., 2014). These outgroup sequences provide the possibility of comparing published mitochondrial genomes for *Turbo cornutus* (National Center for Biotechnology Information [NCBI] GenBank accession NC_061024.1) and *Lunella* aff. *cinerea* (KF700096.1) with the results from our pipeline using the same raw sequence data. Specifically, a blast search was implemented to compare sequence homology. In addition, sequence depth variation, GC content and repeat content were visualised using custom Circos plots (Krzywinski et al., 2009; https://github.com/o-william-white/circos_plot_organelle; accessed 08/2023).

### Human contamination

Prior to running the pipelines, the extent of human contamination was evaluated, a common contaminant in museum samples. Raw reads were adapter trimmed and quality filtered with fastp (S. Chen et al., 2018) and mapped to the human reference genome (NCBI GRCh38.p14) using bwa mem (Li, 2013) and the number of mapped reads quantified with samtools (Li et al., 2009).

### Pipeline descriptions

#### skim2mt

As input, the *skim2mt* pipeline requires two files to be provided by the user: (1) a config file (in YAML format) and (2) a sample list file (in CSV format). The config file outlines the main parameters including the GetOrganelle reference method, adapter sequences used, reference databases used for annotation, alignment trimming methods and outgroup samples for the phylogenetic analyses. The samples.csv file is a list of the samples included in the analysis with the sample names, paths to forward and reverse reads, NCBI taxonomy IDs for searches of reference sequences on NCBI or paths to manually generated gene and seed databases required by GetOrganelle. Note that example config.yaml and samples.csv files are provided with all pipelines (see data and code availability statement). The pipeline accepts NGS data from short read platforms (e.g., Illumina) in paired fastq format.

The *skim2mt* pipeline (Figure 2) starts by processing the data from each sample using fastp (S. Chen et al., 2018) to detect and remove adapter sequences, trim low-quality sequences with parameters for forward and reverse adapter sequences and optionally duplication (`--dedup`) specified. Fastqc is implemented on raw and quality filtered reads (Andrews, 2010). GetOrganelle (Jin et al., 2020) is then used to assemble the target sequence of interest, using seed and gene reference databases to identify and assemble target reads. Although GetOrganelle is provided with default seed and gene databases, our initial benchmarking highlighted that using custom reference databases from closely related taxa minimised the likelihood of assembling contaminant sequences. Therefore, the pipeline will either generate a seed and gene database using the python script go_fetch.py version 1.0.0 (https://github.com/o-william-white/go_fetch), or it will use custom databases provided by the user. The go_fetch.py script takes a NCBI taxonomy ID provided by the user in the samples.csv file, searches the NCBI nucleotide database for mitochondrial reference sequences that are as close as possible to the target taxonomy by working back up the NCBI taxonomy hierarchy until sufficient references are found (minimum 5, maximum 10), and then downloads and formats the reference data for use with GetOrganelle. GetOrganelle is implemented with the following additional parameters: `--reduce- reads-for-coverage inf –max-reads inf -R 20`. Sequences assembled by GetOrganelle are typically named based on the output of SPAdes (Prjibelski et al., 2020), which can produce long sequence names. Therefore, sequences are renamed to <SAMPLE_NAME>_contig <N> if there are multiple contigs or <SAMPLE_NAME>_circular if a single circular sequence is found. Note that GetOrganelle can produce more than one assembled sequence where there are different possible paths through the same assembly graph e.g., mitochondrial genomes containing repeats. However, the pipeline simply selects the first assembled sequence for downstream analyses as the main outputs are the annotated gene sequences and the correct orientation of repeat regions is not necessary. Note that GetOrganelle also generates an assembly graph (.GFA) which can be viewed with bioinformatic tools including Bandage (Wick et al., 2015). Basic assembly statistics are summarised using SeqKit (Shen et al., 2016).

**Figure 1.**
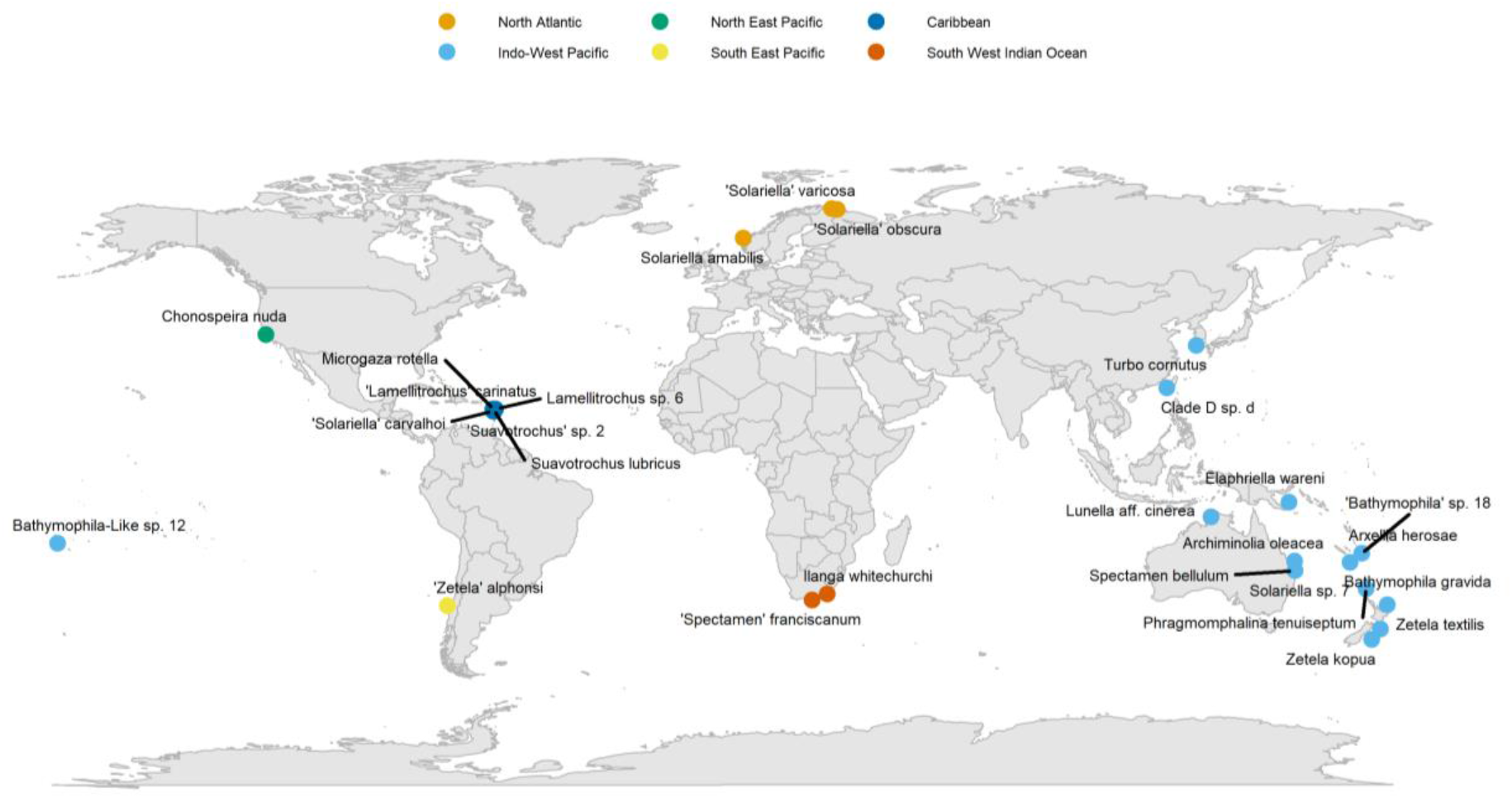
Map showing collection localities for solariellid gastropods samples used in this study.

**Figure 2.**
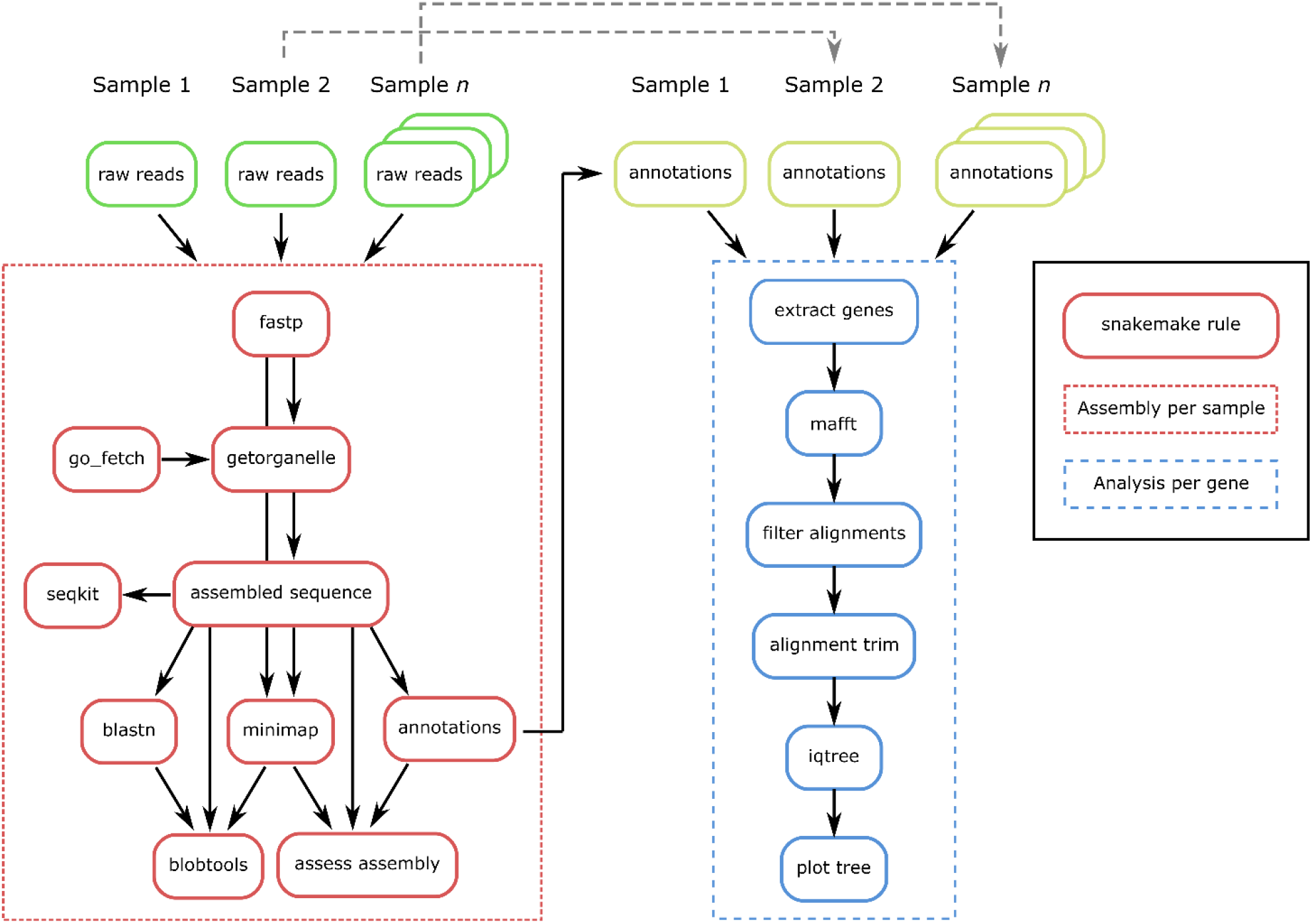
Schematic diagram of *skim2mt* and *skim2rrna*. Both pipelines initially process raw read data from individual samples, assembling target sequences and assessing assembly quality. Each pipeline then aligns, processes, and implements phylogenetic analysis for all annotated genes found across assembled sequences. The *skim2mt* and *skim2rrna* pipelines follow a similar workflow, except for GetOrganelle parameters, and homology search and annotation reference databases.

Next, the assembly quality is evaluated using a blastn search (Camacho et al., 2009) against a temporary blast database created as part of the workflow. For mitochondrial sequences, a blast database generated from the NCBI mitochondrion RefSeq database is used (https://ftp.ncbi.nlm.nih.gov/refseq/release/mitochondrion/). Quality filtered reads generated by fastp are mapped to the assembled sequence using minimap2 to estimate sequence coverage (Li, 2018). The blast and mapping outputs are summarised using blobtools (Laetsch et al., 2017) and the likely taxonomy of the assembled sequence is defined using the taxrule “bestsumorder”. Following the assembly quality check, assembled sequences are annotated using MITOS2 (Bernt et al., 2013). Following assembly and annotation, a plot is created using a custom python script to visualise the location of annotated genes, coverage, and proportion of mismatches in mapped reads.

Once the sequences are assembled and annotated, the checkpoint function of snakemake is used to recover all annotated gene sequences assembled across samples (Figure 2). For each annotated gene recovered, mafft (Katoh & Standley, 2013) is used to align sequences with the following parameters: `--maxiterate 1000 ---globalpair --adjustdirectionaccurately`. To avoid the inclusion of annotated gene sequences with largely missing data in the alignment, individual sample sequences with ≥50% missing data relative to the alignment are removed. Note that the threshold for missing data can be adjusted by the user in the config.yaml input, which may be of particular value when working with very rare specimens. The alignments are trimmed using either Gblocks (Castresana, 2000) or Clipkit (Steenwyk et al., 2020) as specified in the config.yaml file. Phylogenetic analysis is then implemented with IQ-TREE 2 (Minh et al., 2020) using 1000 ultrafast bootstraps and consensus trees are plotted in R using the ggtree package (R Core team, 2020; Yu et al., 2017). Note that phylogenetic analysis is only implemented if there are at least five sequences in the alignment.

#### skim2rrna

The *skim2rrna* pipeline (Figure 2) requires the same input data and follows similar steps to the *skim2mt* pipeline described above, except for the parameters and tools used for the assembly, homology search and annotation. Specifically, for the assembly with GetOrganelle, the following parameters are used: `-F anonym –reduce-reads-for-coverage inf –max-reads inf -R 10 –max-extending-len 100 -P 0`. For the homology search with blast, a blast database generated from the SILVA 138 database is used (Quast et al., 2013). Finally, barrnap (https://github.com/tseemann/barrnap) is used for annotation of ribosomal sequences.

#### Assessing assembly and annotation results

After running *skim2mt* and/or *skim2rrna*, it is necessary to check the results for evidence of contamination across samples or individual genes. This is especially important for museum samples. Specifically, the blobtools (Laetsch et al., 2017) output should be checked for sequences with unusual (e.g. taxonomically divergent) blast hits and the gene alignments and gene trees reviewed by taxonomic experts to identify incongruent relationships. Users may check the summary counts of genes recovered across samples, which can be useful for identifying samples with large amounts of missing data for downstream analyses. Putative contaminant sequences, genes with large amounts of missing sequences in the alignment or samples with large amounts of missing genes can then be removed using the supplementary python script *format_alignments.py* which removes sequences from alignments based on sequence names and formats alignments for use with *gene2phylo*.

#### gene2phylo

After running *skim2mt* and *skim2rrna* and removing putative contaminants, the user may wish to implement further phylogenetic analyses of the filtered alignments. To assist with this, a final pipeline *gene2phylo* is provided to reanalyse assembled genes (Figure 3). As input, the *gene2phylo* pipeline only requires a config file to specify the main parameters for the phylogenetic analysis. The *gene2phylo* pipeline accepts input data from multiple individual genes (aligned or unaligned) in a single directory. The alignments must be named after the gene names (e.g. “cox1.fasta”), and the sample names used for sequences across alignments must be consistent. The *gene2phylo* pipeline begins by optionally re-aligning, removing individual sample sequences with ≥50% missing data relative to the alignment, and trimming poorly aligned regions of alignments, using a similar approach to the pipelines described above. Note, this is only necessary if the input alignments were edited for the removal of contaminant sequences, which is likely to influence alignment characteristics. For each input alignment, phylogenetic analysis is implemented with IQ-TREE2 (Minh et al., 2020) using 1000 ultrafast bootstraps. Input alignments are also combined into a partitioned alignment and a partitioned phylogenetic analysis using IQ-TREE 2 is implemented with 1000 ultrafast bootstrap replicates. In addition, individual gene trees are used to infer phylogenetic relationships using astral (Zhang et al., 2018). Phylogenetic trees from each analysis are plotted using the ggtree package (R Core team, 2020; Yu et al., 2017).

**Figure 3.**
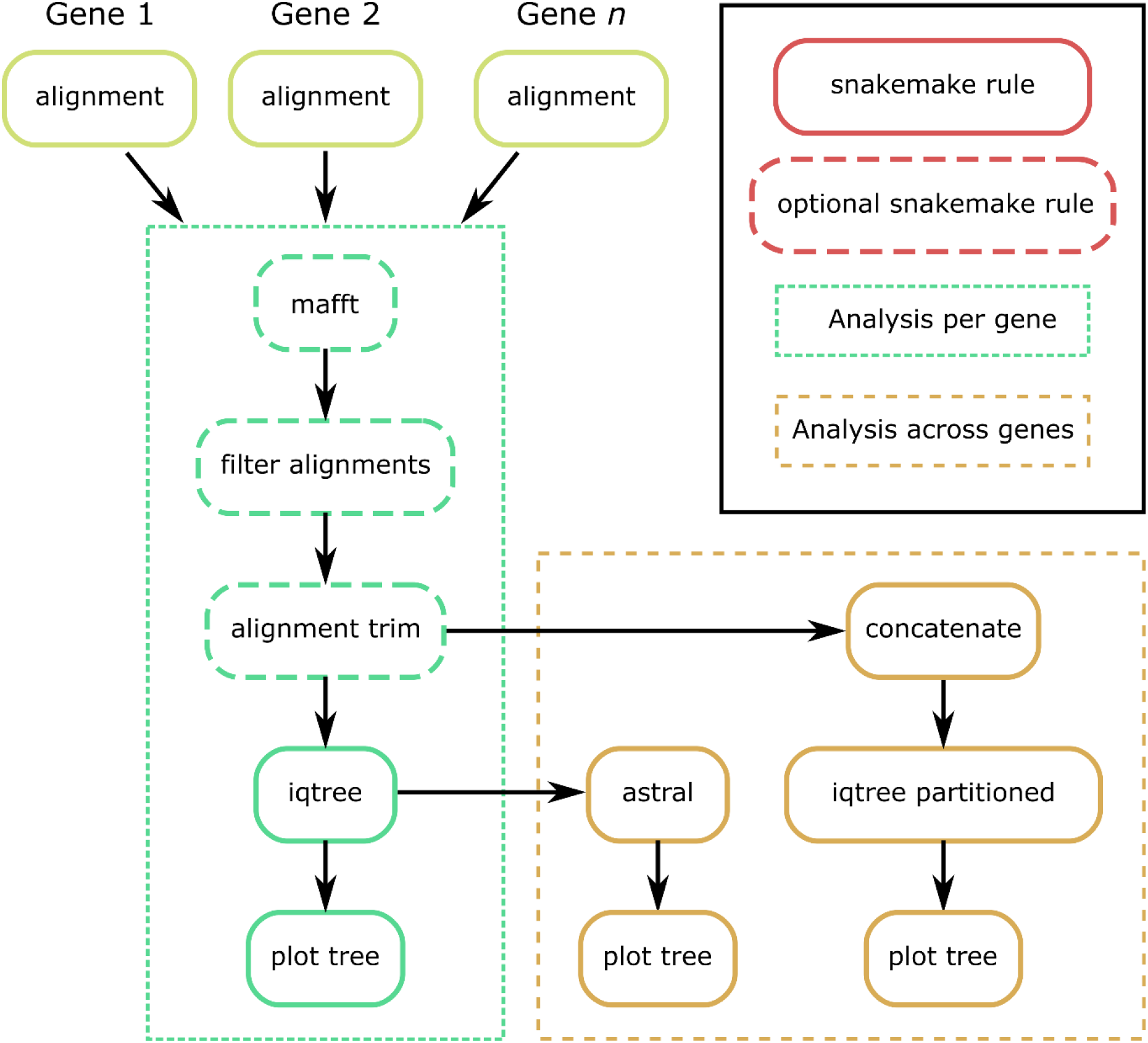
Schematic diagram of *gene2phylo*. Gene2phylo can optionally re-align, remove sequences with too much missing data and trim poorly aligned regions of the alignments. *Gene2phylo* implements IQTREE 2 phylogenetic analysis for each annotated gene, an IQTREE 2 partitioned analysis for all assembled genes and an astral analysis across all individual gene trees.

## Results

### Simulated data

The *skim2mt and skim2rrna* pipeline recovered mitochondrial and ribosomal sequences for all simulated datasets, with a mean assembly size of 15,068 and 6,780 for mitochondrial and ribosomal sequences respectively. The blobtools blast hits predicted the correct taxonomic family for all assembled mitochondrial sequences. However, the correct taxonomic family was only predicted for seven assembled ribosomal sequences. This is likely due to the lack of annotated ribosomal sequences available for the butterfly superfamily Papilionoidea in the SILVA reference data used for the blast database. All annotated genes were retained for phylogenetic analysis with *gene2phylo* and the relationships identified by both the partitioned IQTREE 2 and astral analyses conformed with known subfamily relationships (Supporting information Figure S 1 and Figure S 2).

### Solariellid data

The results of PCR amplification targeting four genes (*cox1*, 28S, 12S and 16S) of our solariellid specimens suggest that many DNA samples were highly degraded (Supporting Information Table S 2). In some cases, faint bands were observed when PCR products were visualised on agarose gels, but clean Sanger sequence could not be obtained, because of low yield and noisy background. DNA quality was confirmed by recording the DNA Integrity Number (DIN) for samples, with a DIN of 10 indicating highly intact DNA fragments, whilst a DIN of 1 indicates a highly degraded DNA sample. DNA quality for the samples used in this study ranged from not detectable for the poorest samples to 6.5 for the best (Supporting Information Table S 2).

Approximately 870 M raw sequence reads were generated across all samples, with an average of 32 M raw reads per sample (Supporting Information Table S 2). On average, 95.62% of reads were retained following adapter removal and basic filtering with fastp. Sequence data contamination with reads originating from human DNA was extensive across samples, with an average of 61.86% (range 32.74%-80.02%; Supporting Information Table S 2).

The *skim2mt* pipeline successfully recovered mitochondrial genome sequences from 24/27 samples with an average assembly size of 14,441 bp (range 347-24,670 bp). A circular mitochondrial genome was assembled for a single sample (*Zetela kopua*; Figure 4). However, no mitochondrial sequences could be assembled for *Elaphriella wareni*, *‘Spectamen’ franciscanum* or *Zetela textilis*. Assembled sequences for outgroups *Turbo cornutus* and *Lunella* aff. *cinerea* were compared to previously published sequences. Blast hits for these two specimens had 100% percentage sequence similarity when compared to previously published sequences, although the sequences assembled by our pipeline were not as complete as those published on GenBank. The assembled sequence for *Turbo cornutus* matched the published sequence (NCBI accession NC_061024.1; Supporting Information Figure S 3) from 1-13,676 and 14,107-17,299. The assembled sequence for *Lunella* aff. *cinerea* matched the published sequence (KF700096.1; Supporting Information Figure S 4) from 1-13,973 and 14,412-17670. Visualisation of circos plots for published circular sequences suggests that the assembly process broke in regions of low coverage, low GC content and/or high repeat content (Supporting Information Figure S 3-4). Of the 15 mitochondrial genes annotated by MITOS2 (13 protein coding genes and two mitochondrial ribosomal subunits), an average of 12 genes were annotated across samples, with 15/27 samples having all protein coding and rRNA genes annotated.

**Figure 4.**
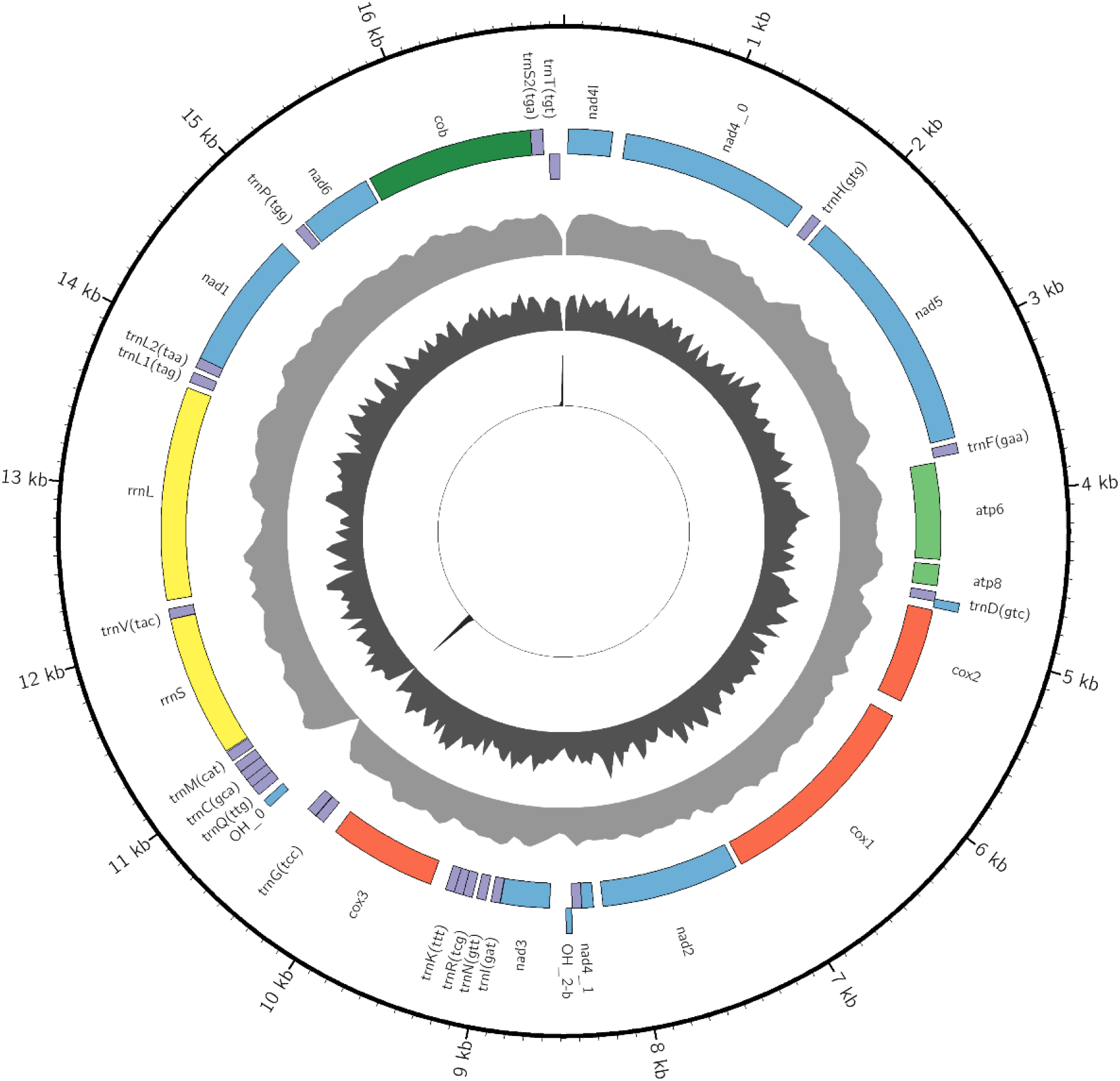
Assembled circular sequence for *Zetela kopua* with the following attributes from outside to inside: sequence position, annotation names, annotations on the + strand, annotations on the – strand, coverage (max=2,779), GC content (max=0.6) and repeat content (max=1.0). This image was created using a custom organelle visualisation tool available on GitHub (https://github.com/o-william-white/circos_plot_organelle; accessed 08/2023).

The *skim2rrna* pipeline successfully recovered ribosomal gene sequences from 27/27 samples with an average size of 3,049 bp. Of the ribosomal genes annotated by barnnap, the 18S, 28S and 5.8S rRNA genes were annotated in 4, 24 and 4 samples respectively.

After checking the outputs of *skim2mt* and *skim2rrna*, 18S and 5.8S rDNA genes were found to have more than 50% missing data across samples and were therefore removed from downstream analyses. In addition, *atp8* was discarded as it was <100 nucleotides in length and contained little phylogenetic information. Six samples were found to be missing more than 50% of the annotated genes and were therefore discarded from all downstream phylogenetic analysis. Excluded species were *Archiminolia oleacea*, *Arxellia herosae*, *Bathymophila gravida*, *Elaphriella wareni*, *‘Spectamen’ franciscanum*, and *Zetela textilis*. After manual checking of the blobtools output, alignments and phylogenetic trees generated by *skim2mt* and *skim2rrna*, it was determined that although all genes were annotated for *Spectamen* cf. *bellulum,* these data likely originated from another solariellid species as a contaminant, from the genus *Ilanga*. Annotations for 28S from *Phragmomphalina tenuiseptum* and *Zetela kopua* were also identified as likely non-solariellid gastropod contaminants. Duplicate gene annotations were identified in Clade D sp. d (*cox3*, *nad3*, *nad2*, *cox1*, *cox2*, *atp8*, *atp6*), *Solariella amabilis* (*nad3*) and *Zetela alphonsi* (nad2). Where duplicate genes were identified, only the first annotation was used in downstream analyses. Following the removal of genes and samples with too much missing data, contaminant sequences and duplicate annotations, the final dataset consisted of 15 genes for 20 specimens with an average 6.33% missing data on average across samples.

With the filtered set of genes, *gene2phylo* was implemented to re-align and analyse the individual gene alignments. Phylogenetic analysis of the partitioned alignment in IQ-TREE (Figure 5) recovered a tree with support values ranging from poor to optimal (29-100%). Phylogenetic analysis using the individual genes trees with astral (Supporting Information Figure S 2) recovered a tree with broadly consistent topology.

**Figure 5.**
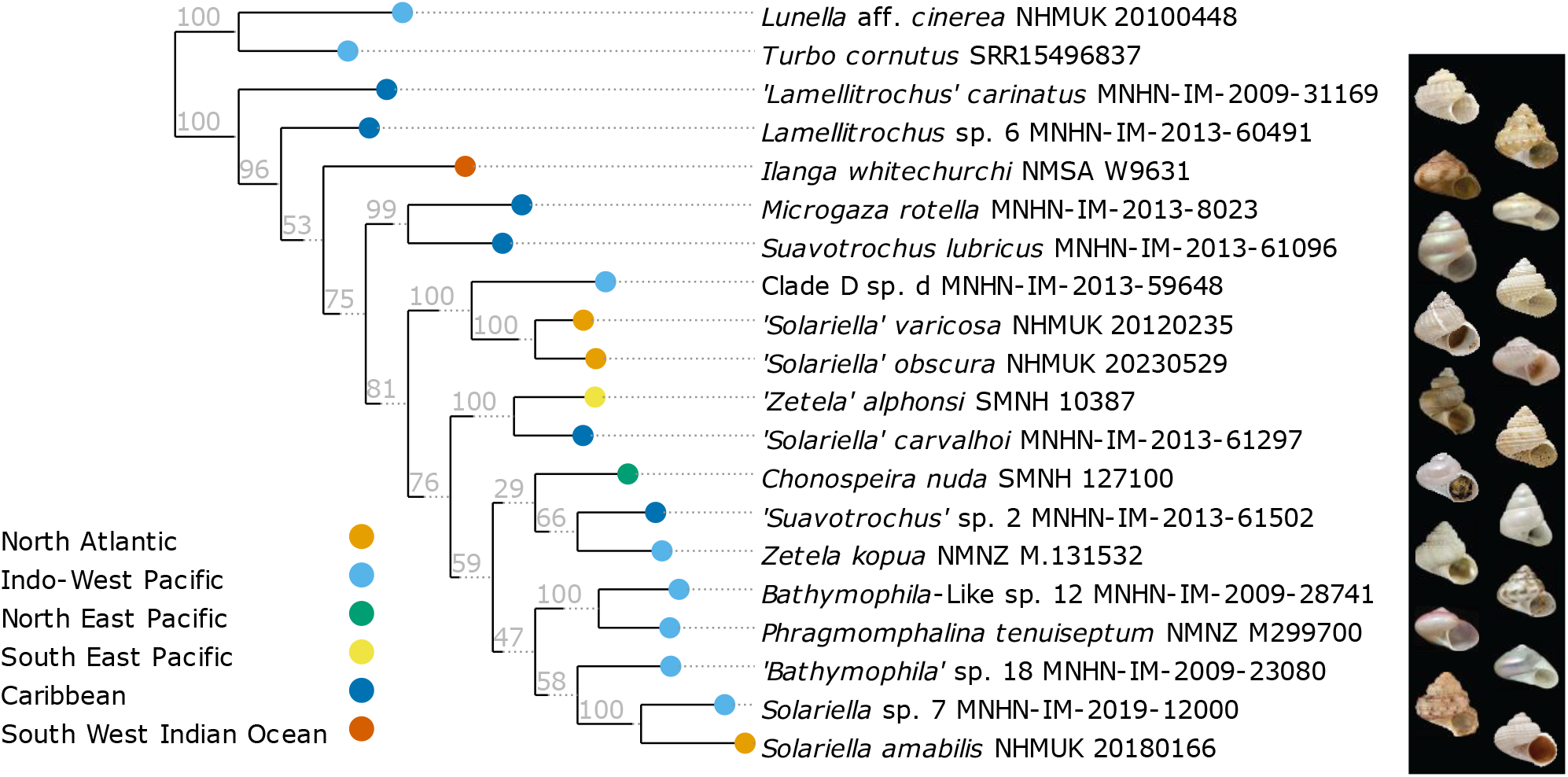
Partitioned maximum likelihood tree of 15 genes including 12 mitochondrial protein-coding genes, two mitochondrial ribosomal genes and one nuclear ribosomal gene (28S) generated using IQ-TREE 2 and visualised using ete3. The tree is rooted on the outgroup taxa and values on branches are ultrafast bootstrap values. Images of each specimen including in the analysis are provided, except for outgroups *Lunella aff. cinerea* and *Turbo cornutus*.

## Discussion

This study demonstrates the utility of a snakemake toolkit comprised of the pipelines *skim2mt, skim2rrna* and *gene2phylo*, for the assembly, annotation, and phylogenetic analysis of mitochondrial and ribosomal genes from genome skimming datasets. Analysis of simulated data from the butterfly superfamily Papilionoidea is used to benchmark our pipelines, generating expected phylogenetic relationships. Further analysis of novel sequence data from the gastropod family Solariellidae, generated some of the first mitochondrial genomes and ribosomal sequences for the family. Specifically, complete or partial mitochondrial genomes were obtained for 23/27 non-contaminated specimens, and ribosomal sequences were assembled for 24/27 samples.

The Solariellidae samples included in this study represent many of the issues that are typical of historical museum specimens. For example, three samples (*‘Solariella’ obscura*, *‘Solariella’ varicosa* and *Solariella amabilis*) were collected more than 50 years ago and preserved in low percentage ethanol (70%) with uncracked shells. In addition, DNA was extracted more than ten years ago from dehydrated tissue samples of another sample (*Bathymophila*-Like sp. 12). Therefore, several of the Solariellidae samples had highly degraded DNA (DIN<2). Despite this, our pipelines were able to recover nearly complete mitochondrial genomes and ribosomal sequences for most samples.

Contamination is a common issue with historical specimens. Indeed, human contamination was extensive across all the Solariellidae samples sequenced in this study, with an average of 61.86% human reads per sample. This is likely due to the small amount of degraded DNA from the target specimen and the coextraction of human DNA from contact with samples during collection, curation, tissue sampling and laboratory work. Despite this, our pipelines avoided the assembly of human sequences by using reference databases from closely related species. The assembly of contaminant sequences from closely related species is more problematic, but these contaminants can be identified by sequence homology searches and phylogenetic analysis. From our analysis, we were able to determine that all sequences obtained for *Spectamen* cf. *bellulum* are likely contaminants from a morphologically distinct solariellid sequenced previously in the same lab. In addition, 28S annotations from *Phragmomphalina tenuiseptum* and *Zetela kopua* were also identified as likely contaminants from non-solariellid gastropods, also sequenced in the same lab. Therefore, these contaminants are likely due to laboratory contamination, which can be minimised by switching from a standard bench to clean room facilities and following protocols used for ancient DNA extractions.

The mitochondrial sequences assembled for outgroups *Turbo cornutus* and *Lunella* aff. *cinerea* showed high sequence homology to previously published sequences, although the assemblies were not as complete as those published on GenBank, with assemblies appearing to break in regions of low coverage, low GC content and/or high repeat content (Supporting Information Figure S 3 ∼13,676 bp; Figure S 2 ∼13,973 bp). Using a reference-based organelle assembly tool such as MITObim (Hahn et al., 2013) may increase the likelihood of a complete assembly, but at the risk of reference bias. However, for studies such as this, selecting a single reference may be problematic when working with a diverse range of taxa, with complex variation in organelle genome sequences. Our pipeline utilises GetOrganelle which uses a *de novo* assembly approach and so should have a smaller reference bias, even from a diverse range of taxa. In addition, the most important output for the phylogenetic analysis is the list of annotated genes used in phylogenetic analyses, which were completed for both our outgroup samples.

Genome-skimming offers many advantages over traditional PCR amplification. Indeed, genome skimming and the *skim2mt and skim2rrna* pipelines were able to recover gene sequences that were previously unattainable by PCR. For example, *cox1* could be sequenced using Sanger sequencing for only 8/25 samples (not including outgroups), whereas *cox1* was recovered for 18/25 samples using genome skimming and *skim2mt*. Likewise, 28S could only be sequenced from PCR amplicons for 9/25 samples whereas genome skimming assembled 28S for 23/25 samples. The latter is likely due to DNA degradation since primer sequences come from a very conserved region, whereas some *cox1* failures may also be due to primer mismatches.

Previous phylogenetic analyses of solariellid gastropods have highlighted complex and unresolved phylogenetic relationships among genera (Williams et al., 2020). Although the tree in this study has good (>95%) ultrafast bootstrap support for most terminal clades, support for basal splits often have poorer support, suggesting that some assignments to genera require further research. One potential option would be to increase taxon sampling to test generic assignments. Likewise, while individual gene sequences from barcode genes are already available for a diverse range of solariellid gastropods, a greater sampling of gene space is required to confirm the relationships within this group.

Although this methodology will work for any sample type, *skim2mt and skim2rrna* were written specifically to account for many of the issues associated with historical museum samples, for example, DNA degradation and contamination. By default, *skim2mt* implements GetOrganelle using all reads as input (`--reduce-reads- for-coverage inf --max-reads inf`) and an increased number of rounds of (`-R 20`) of target read selection. For museum samples that are likely to be degraded, this maximises the inclusion of short sequencing reads. In addition, the user can specify a custom reference database for GetOrganelle using sequences from closely related taxa. This is necessary because our benchmarking of GetOrganelle using simulated datasets (White and Clark unpub. data) highlighted that a reference dataset containing closely related sequences increases the likelihood of successful assembly. Conversely, a broad reference dataset can increase the likelihood of sequence assembly from contaminated DNA. Taxonomic assignment of assembled sequences using blobtools can identify most non-target sequences, and phylogenetic analysis implemented for all annotated gene sequences may be particularly useful for identifying genetically similar contaminants not recognised by blobtools.

Although the pipelines presented simplify the bioinformatic analyses significantly, allowing for the analysis of hundreds of samples simultaneously, there is risk to trade-off accuracy for increasing scale of the analysis. Indeed, our study highlighted that it was still important to manually check the assembled sequences for contamination or poorly annotated sequences using the standard outputs of our pipeline including blobtools, individual gene alignments and phylogenetic analyses. Although, it could be hardcoded to remove assembled sequences based on sequence homology to reference databases, this is not possible at present, because public databases are not complete for all taxa. In addition, taxonomic expertise may be necessary to identify incongruent phylogenetic relationships that can result from cross contamination from closely related taxa, highlighting the need for particular care when extracting DNA from historical specimens. An additional pipeline could be further adapted to include chloroplast organelle sequences by including a chloroplast annotation tool. However, there are few chloroplast annotation resources available as a command line tool with premade reference databases that annotate genes in a consistent and reliable way, as MITOS2 does for mitochondrial genes.

In conclusion, this study demonstrates that the snakemake pipelines *skim2mt*, *skim2rrna* and *gene2phylo* can cope with poor quality data from historical collections, facilitating large scale genome skimming studies from museum specimens. Given the current biodiversity crisis and lack of taxonomic expertise, it has become more important than ever to document biodiversity before it is lost. By sequencing natural history collections at scale using bioinformatic tools such as those presented here, researchers can increase the rate of phylogenetic and barcoding studies, and ultimately, species discovery.

## Acknowledgements

We thank Andrea Waeschenbach for sequence data for *‘Solariella’ varicosa;* QuantaBio for providing the sparQ DNA Library Prep kit and sparQ PureMag Beads; Tore Høisaeter for providing specimens of ‘*Solariella’ obscura* and ‘*Solariella’ varicosa*; Anders Warén for providing specimens of *Solariella amabilis* and loaning specimens of *Chonospeira nuda* and ‘*Zetela’ alphonsi*; Dai Herbert for loaning specimens of *Ilanga whitechurchi* and ‘*Spectamen’ franciscanum*, Ian Loch for loaning specimens of *Archiminolia oleacea*; Bruce Marshall for loaning *Bathymophila gravida*, *Zetela textilis, Z. kopua* and the holotype of *Phragmomphalina tenuiseptum* and Barbara Buge, Nico Puillandre and Philippe Bouchet for loaning the remaining specimens from MNHN. Thanks also to Jon Kongsrud for specimen locality data and to Harry Taylor for photos of NHMUK and MNHN specimens, Bruce Marshall for the photo of *Zetela kopua* and Sadie Mills for the photo of *Phragmomphalina tenuiseptum*. MNHN Specimens were obtained during research cruises and expeditions organized by the MNHN with Institut de Recherche pour le Développement and National Taiwan University, as part of the Tropical Deep-Sea Benthos program: ZhongSha 2015 in the South China Sea; MADEEP (dx.doi.org/10.17600/14004000) in Papua New Guinea; BERYX 11 (dx.doi.org/10.17600/92005011), EXBODI (dx.doi.org/10.17600/11100080) and NORFOLK 1 (dx.doi.org/10.17600/1100050) in New Caledonia; BORDAU 1 (dx.doi.org/10.17600/99100020) in Fiji; KARUBENTHOS 2012 and KARUBENTHOS 2 (dx.doi.org/10.17600/15005400) in Guadeloupe (more information can be found at expeditions.mnhn.fr). These expeditions operated under the regulations then in force in the countries in question and satisfy the conditions set by the Nagoya Protocol for access to genetic resources. Finally, we would like to thank Rutger Vos and Dick Groenenberg for their initial feedback on the pipelines.

## Data and code availability statement

Raw sequence reads can be downloaded from the NCBI short read archive under BioProject (to be submitted upon manuscript submission), with the exception of previously published data for *Turbo cornutus* (SRR15496837). The snakemake pipelines are available on both GitHub and WorkflowHub: skim2mt https://github.com/o-william-white/skim2mt, skim2rrna https://github.com/o-william-white/skim2rrna and gene2phylo https://github.com/o-william-white/gene2phylo. Note that each pipeline is provided with the simulated data from 25 species of the Butterfly superfamily Papilionoidea used in the manuscript, with instructions on how to run each pipeline. Configuration files used for the Solariellidae analyses are available on Data Dryad (To be completed upon manuscript submission). Assembled mitochondrial genome sequences except for previously published outgroups are available from the NCBI nucleotide database under the following accession records: (To be completed upon manuscript submission).

## Supporting Information

### Tables

**Table S 1 Simulated taxa and reference mitochondrial and ribosomal sequences used.**

**Table S 2 Summary of sample quality for extracted DNA and details of factors affecting DNA including the year of sample collection, preservative (ethanol or dry shell), if shell was cracked to allow penetration of ethanol, year DNA was extracted, DNA Integrity Number (DIN) and the amplification success for four partial gene sequences (mitochondrial genes: cox1, 16S rRNA and 12S rRNA, nuclear 28S rRNA). PCR success is summarised as GenBank number if PCR were successful and published, "SEQ" if the PCR was successful and sequenced but unpublished, or "PCR only" if a band was observed when the PCR amplicon was run on a gel but attempts to sequence the amplicon were unsuccessful. Raw and quality filtered sequencing reads generated by fastp are summarised. Mitochondrial genome and ribosomal gene assembly is summarised as assembled base pairs and mapped reads. Notes on the sample filtering implemented prior to the final phylogenetic analysis are presented. Finally, a count of mitochondrial and ribosomal genes recovered are shown.**

### Figures

**Figure S 1.**
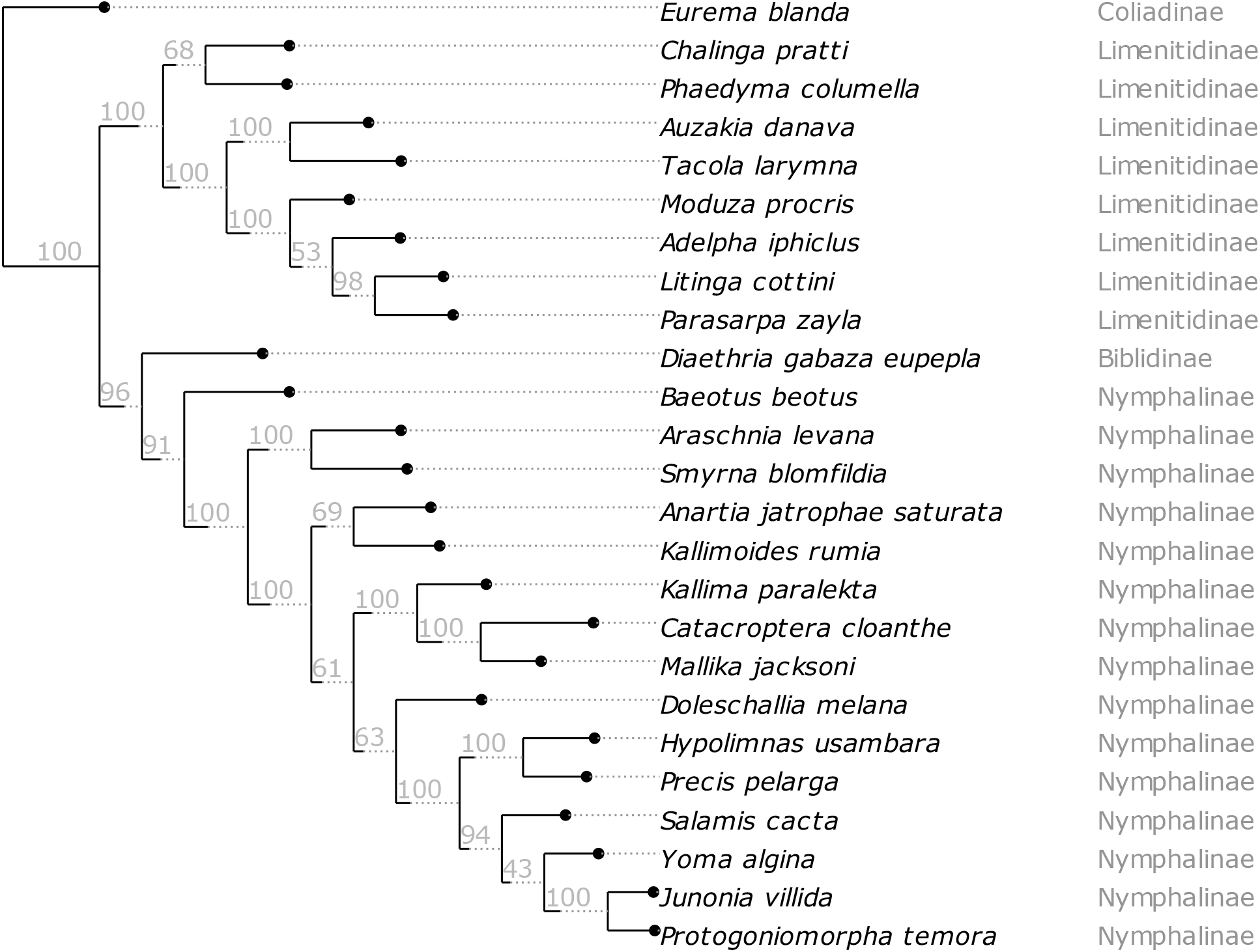
Partitioned IQTREE 2 from analysis of simulated data from 25 published organelle and ribosomal genomes. The analysis included 18 genes including 13 mitochondrial protein-coding genes, two mitochondrial ribosomal genes and three nuclear ribosomal genes and visualised using ete3. The tree is rooted on the outgroup taxon *Eurema blanda* and values on branches are ultrafast bootstrap values. Tips are labelled with species and subfamily names.

**Figure S 2.**
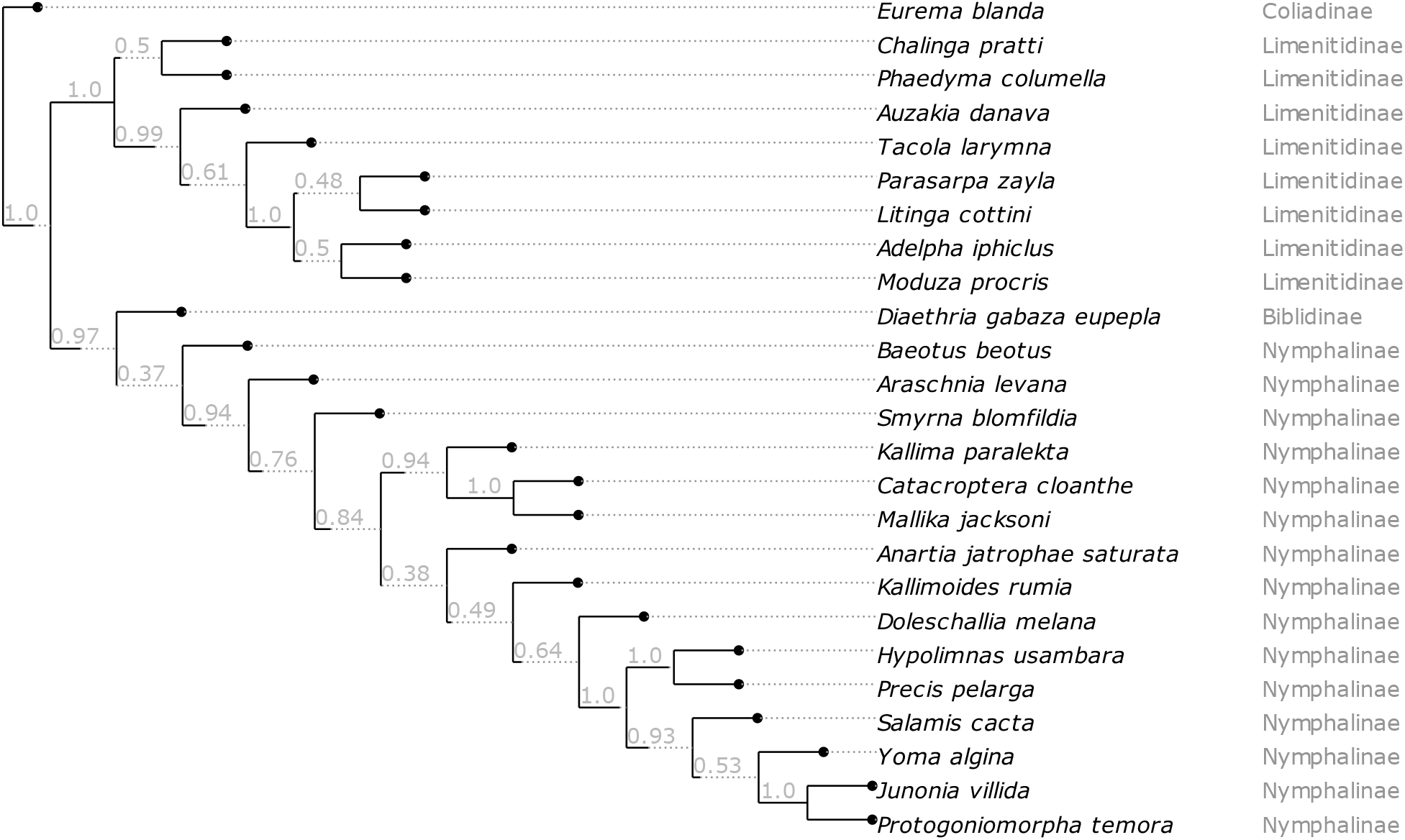
Astral tree from analysis of simulated data from 25 published organelle and ribosomal genomes. The analysis included 18 genes including 13 mitochondrial protein-coding genes, two mitochondrial ribosomal genes and three nuclear ribosomal genes visualised using ete3. The tree is rooted on the outgroup taxon *Eurema blanda* and values on branches are bootstrap values. Tips are labelled with species and subfamily names.

**Figure S 3.**
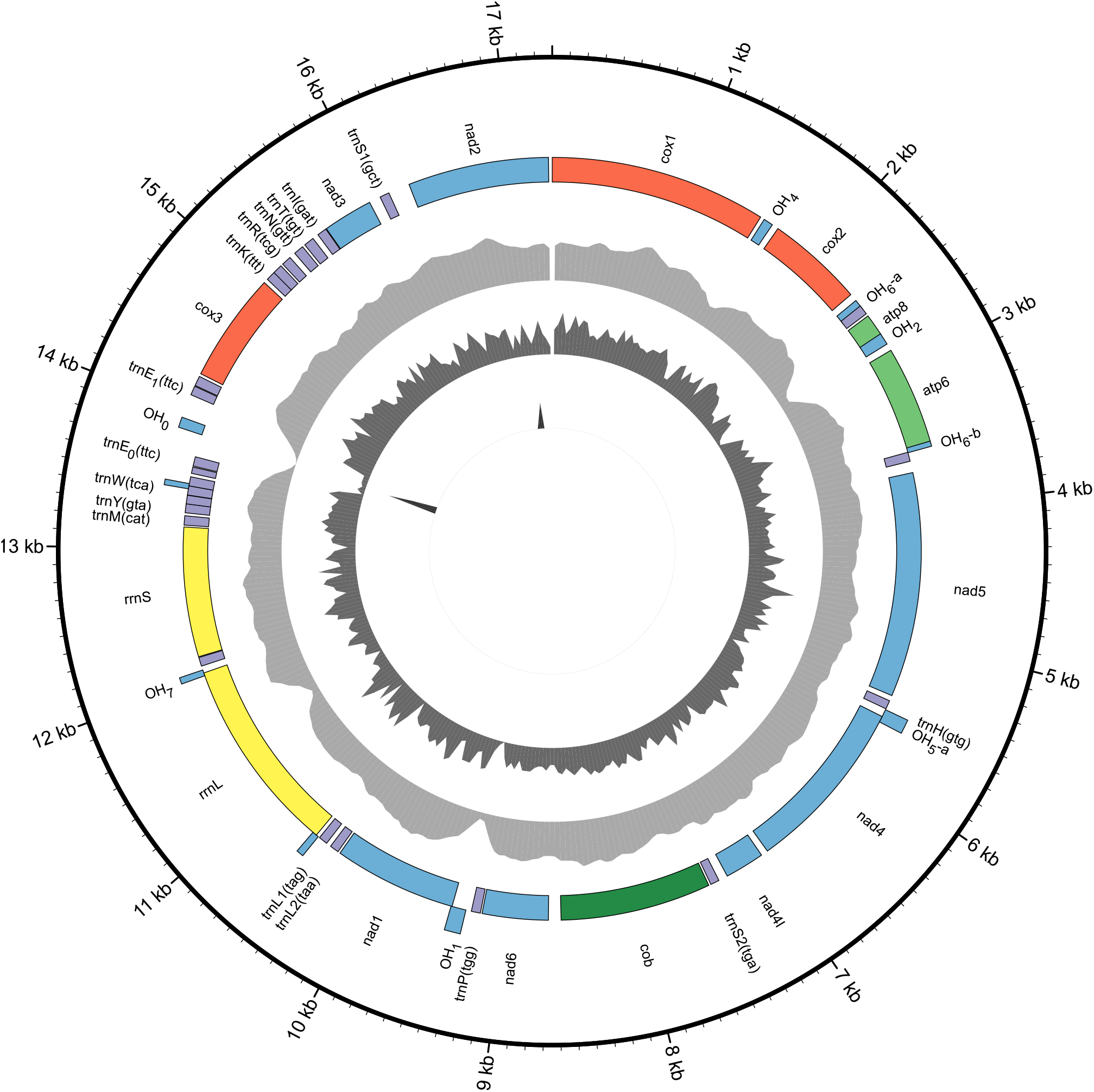
Previously published assembled sequence for *Turbo cornutus* (NCBI accession NC_061024.1) with the following attributes from outside to inside: sequence position, annotation names, annotations on the + strand, annotations on the - strand, coverage (max=2779), GC content (max=0.6) and repeat content (max=1.0). This image was created using a custom organelle visualisation tool available on GitHub (https://github.com/o-william-white/circos_plot_organelle; accessed 08/2023).

**Figure S 4.**
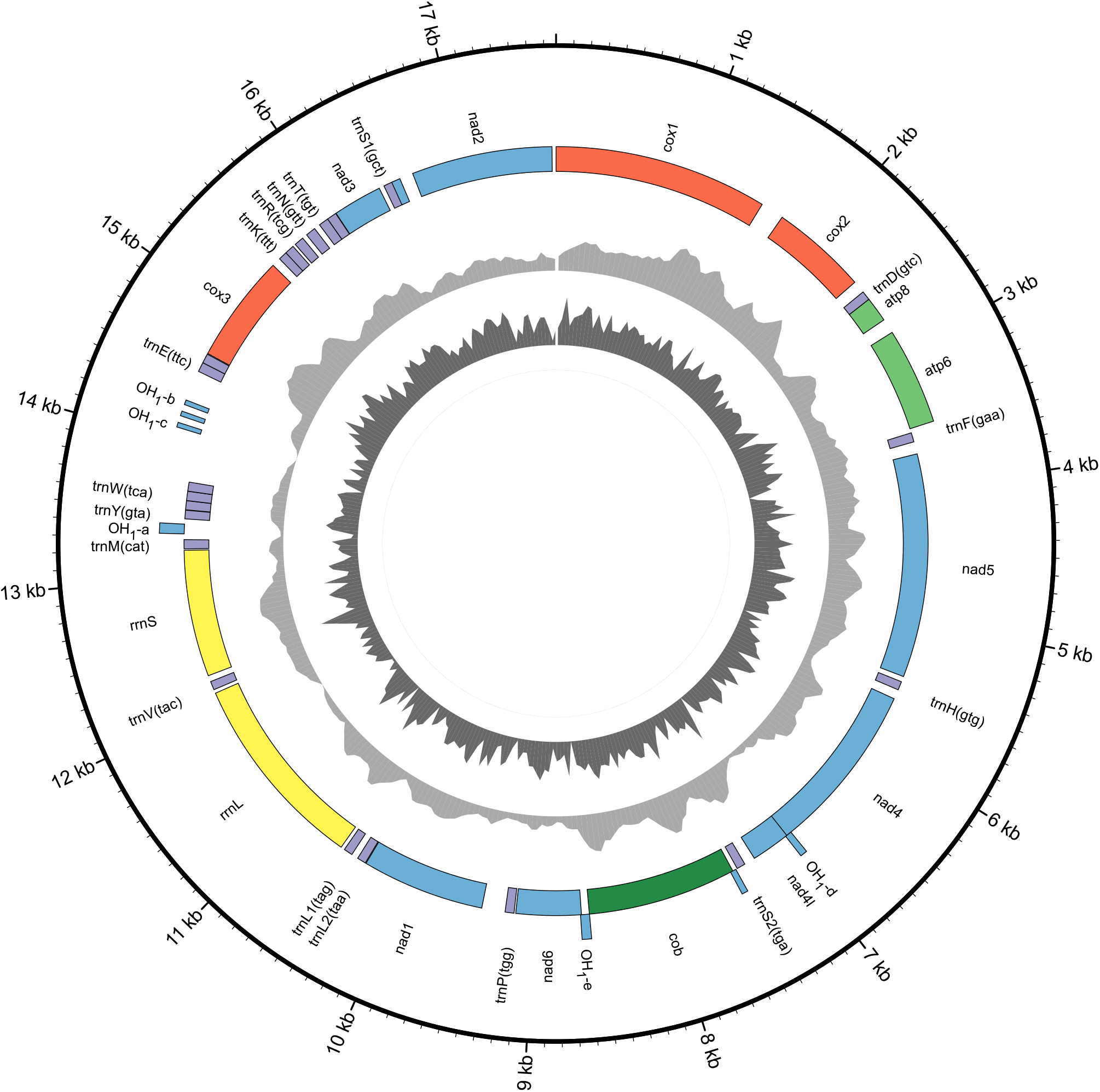
Previously published assembled sequence for *Lunella* aff. *cinerea* (NCBI accession KF700096.1) with the following attributes from outside to inside: sequence position, annotation names, annotations on the + strand, annotations on the - strand, coverage (max=2779), GC content (max=0.6) and repeat content (max=1.0). This image was created using a custom organelle visualisation tool available on GitHub (https://github.com/o-william-white/circos_plot_organelle; accessed 08/2023).

**Figure S 5.**
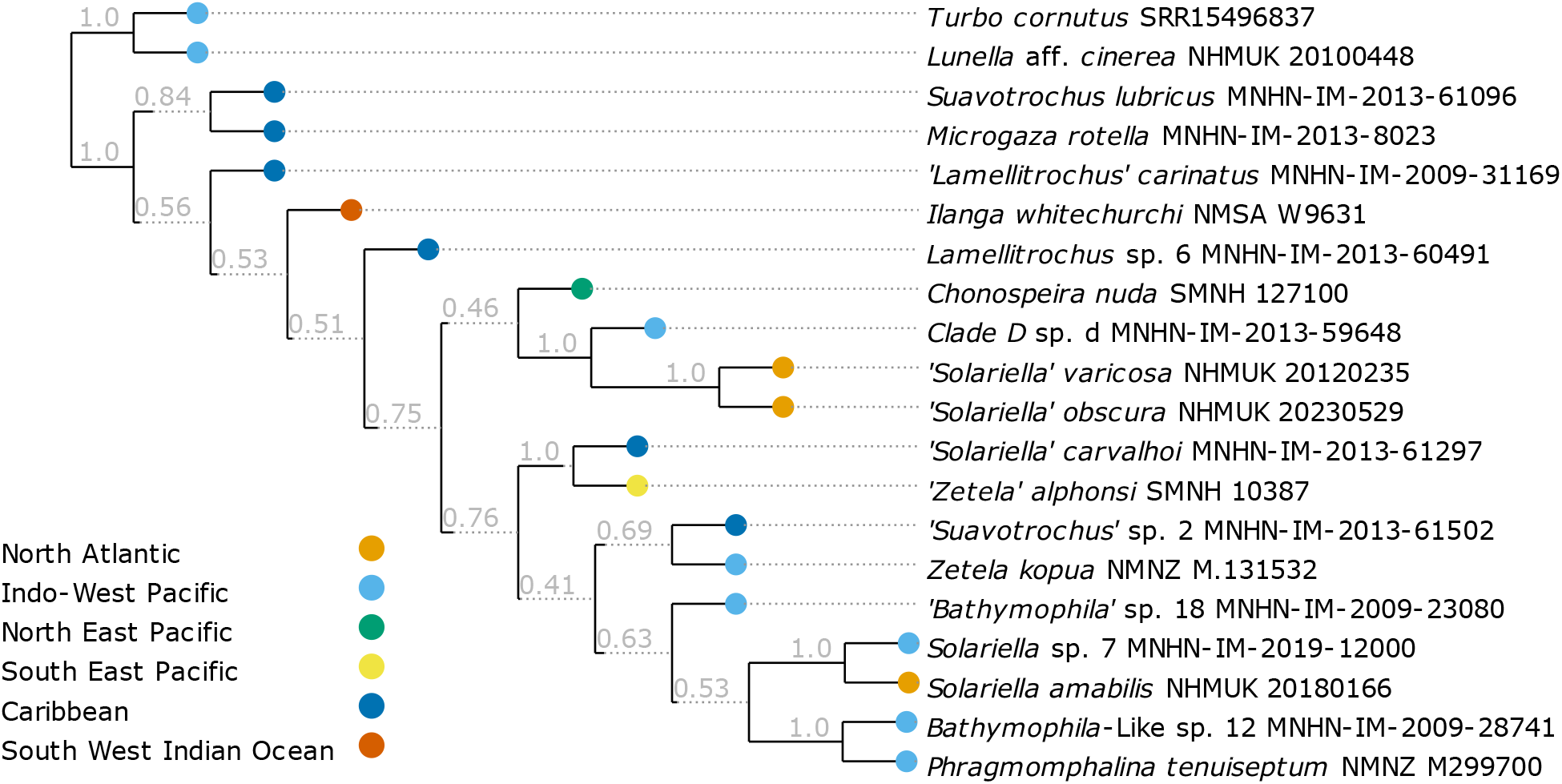
Astral tree of 15 genes including 12 mitochondrial protein-coding genes, two mitochondrial ribosomal genes and one nuclear ribosomal gene (28S) and visualised using ete3. The tree is rooted on the outgroup taxa and values on branches are bootstrap values.

